# Influence of data sampling methods on the representation of neural spiking activity *in vivo*

**DOI:** 10.1101/2022.01.11.475844

**Authors:** Meike E. van der Heijden, Amanda M. Brown, Roy V. Sillitoe

## Abstract

*In vivo* single-unit recordings distinguish the basal spiking properties of neurons in different experimental settings and disease states. Here, we examined over 300 spike trains recorded from Purkinje cells and cerebellar nuclei neurons to test whether data sampling approaches influence the extraction of rich descriptors of firing properties. Our analyses included neurons recorded in awake and anesthetized control mice, as well as disease models of ataxia, dystonia, and tremor. We find that recording duration circumscribes overall representations of firing rate and pattern. Notably, shorter recording durations skew estimates for global firing rate variability towards lower values. We also find that only some populations of neurons in the same mouse are more similar to each other than to neurons recorded in different mice. These data reveal that recording duration and approach are primary considerations when interpreting task-independent single-neuron firing properties. If not accounted for, group differences may be concealed or exaggerated.

## Introduction

*In vivo* recordings of single neurons are used to examine and compare spiking activity between distinct cell types ^1–3^, brain regions ^4^, developmental timepoints ^5,6^, disease models ^7,8^, across species ^9^, and among human patients ^10–12^. Many experimental settings render chronic or long-term recordings unfeasible. Experimental constraints, such as when the brain is actively growing, may prevent the implantation of a chronic recording setup ^5,6^. Additionally, transient recordings may be the best or one’s only chance to record specific cell populations, such as opportune recordings of neurons that are obtained in human patients when electrodes are implanted during deep brain stimulation surgery ^10–12^. These shorter-term recordings often measure task-independent or basal firing patterns. Although such recordings have provided fundamental insight into neural function in health and disease, it remains unclear whether short recordings are rich enough to represent the population, and if they are, what are the specific limitations compared to longer recordings. In lieu of chronic recordings or task-specific responsiveness, many experimenters set a predetermined recording duration or spike number as inclusion criterion for analyses of neural firing properties ^4,6,12,13^. Due to the opportunistic nature of these recordings, experimenters must also decide how many neurons and patients or animals to attempt to include in these analyses. Here, we investigate how data sampling strategies may influence the representation of firing properties of single neurons. We analyzed more than 300 experimentally obtained spike trains from two primary populations of cerebellar neurons, Purkinje cells and cerebellar nuclei neurons, which were recorded in control mice as well as in different disease models and behavioral states.

The cerebellum provides unique advantages for determining the impact of sampling strategies on the representation of neuronal firing. One major advantage of the cerebellum is that its circuit is well-defined and has stereotyped connectivity between specific neuronal populations ^14,15^. For example, Purkinje cells, a principal cell type of the cerebellum, integrate incoming motor and sensory information to the cerebellum and form the sole output of the cerebellar cortex. Cerebellar nuclei projection neurons receive major inhibitory input from Purkinje cells and, in turn, form the primary connection between the cerebellum and dozens of other brain regions (Figure 1A, bottom). Purkinje cells and nuclei neurons have intrinsic properties that allow them to generate remarkably regular action potentials even without input from other neurons ^16–18^. However, sensory and motor afferents modify these intrinsic firing properties in the intact cerebellums of live animals ^19–26^, resulting in highly dynamic firing properties in the *in vivo* circuit.

**Figure 1.**
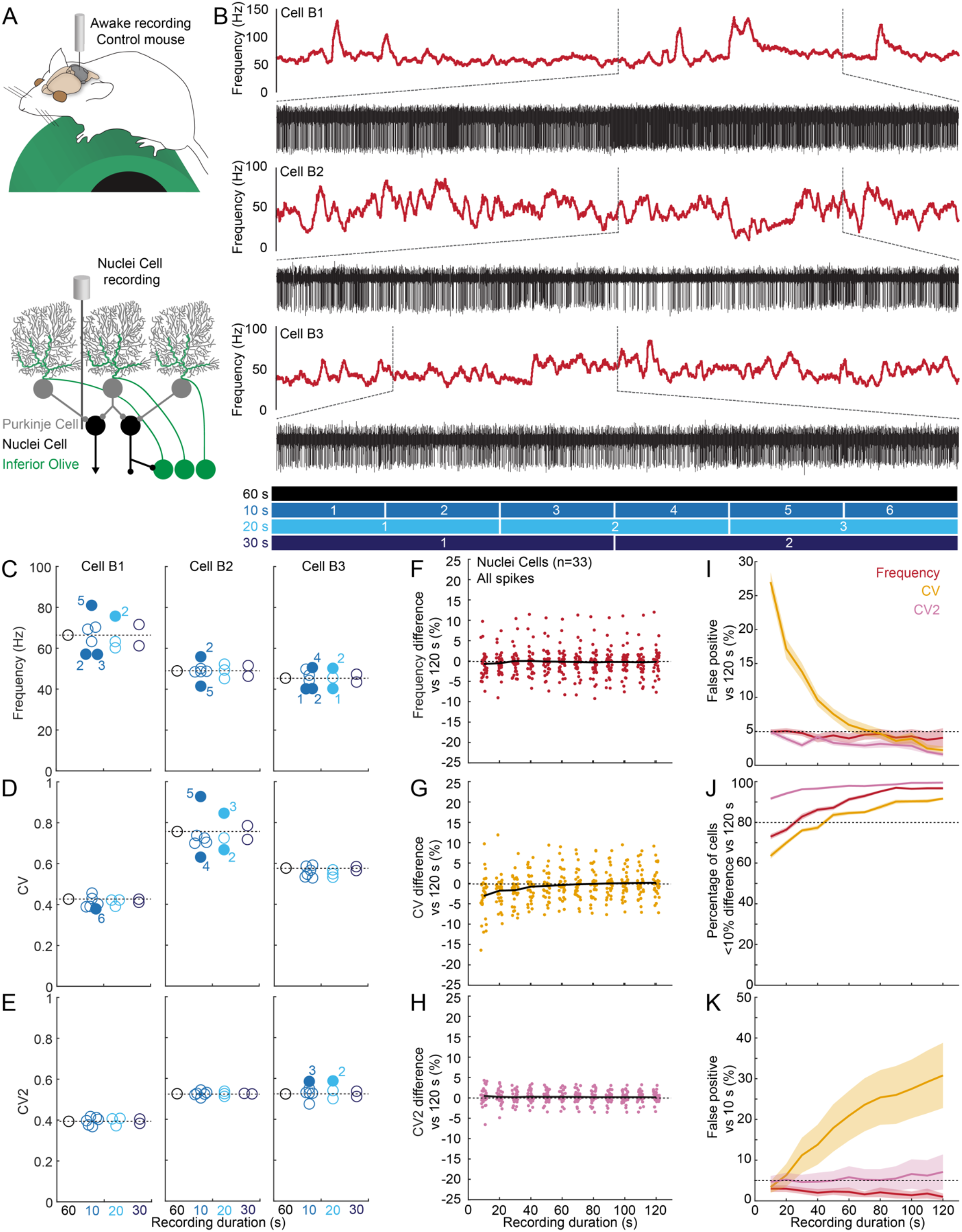
Variability in the firing properties of fast-firing neurons in awake mice. **A**. Schematic of recording setup for awake mice on a foam running wheel (top). For simplicity, in all schematics we have not drawn the plates used for head-fixing the mouse or the recording chamber (see methods for details). Schematic of cerebellar circuit with nuclei neuron recording (bottom). **B**. Three examples of nuclei neuron firing properties. Red traces represent the firing frequency calculated over 0.5 s intervals. The red traces each are 60 s long. The black traces show raw electrophysiological recordings. Each vertical line is an extracellularly recorded action potential. The black traces each are 20 s long and represent the underlying action potentials from the bracketed region of the above red trace. **C**. Frequency, **D**. CV, and **E**. CV2 as calculated for the representative traces in “B.” For C, D, and E: Black circles and dotted lines are the average frequency in the entire 60 s recording. Blue circles are the average frequencies calculated over shorter intervals (for left to right: 10 s, 20 s, and 30 s). Filled circles indicate that the parameter in the designated segment deviates more than 10% from average. Numbers correspond to the segments as indicated in B. **F**. Frequency, **G**. CV, and **H**. CV2 difference from a randomly sampled 120 s recording duration, extracted from a recording of at least 180 s. Each dot represents a single neuron’s average percentage difference of 100 randomly sampled recording durations. The black line represents the average deviation from the frequency, CV, or CV2 calculated from the 120 s-long (reference) recordings. The black line diverging from the dotted line at zero in “G” shows systematic underestimation of CV in shorter recording durations. **I**. Number of significant paired t-tests between 25 randomly sampled 120 s-long (reference) recordings (mean = solid line, +/- SEM = shaded region) and 100 randomly sampled recordings for each varied recording duration for each neuron tested. Frequency is shown in oxblood red, CV in orange, CV2 in pink. **J. P**ercentage of neurons within 10% deviation from the frequency (oxblood red), CV (orange), and CV2 (pink) of a 120 s recording, repeated for 25 randomly sampled 120 s-long (reference) recordings (mean = solid line, +/- SEM = shaded region), each with 100 random samples of each varied recording duration for each neuron. **K**. Number of significant paired t-tests between 100 randomly sampled test duration recordings and 25 randomly sampled 10 s-long (reference) recordings for each cell tested. Mean = solid line, +/- SEM = shaded region.

The modular organization of cerebellar circuits increases the complexity of firing activity within neuron types ^27–29^. Purkinje cell firing frequency and pattern are distinct between at least two populations ^1,2,30,31^, which are consistent with zebrinII molecular expression and specific pre- and post-synaptic partners ^32–34^. The modular organization further influences the firing activity of the cerebellar nuclei projection neurons ^35^, which are classified as excitatory and inhibitory neurons, each with their specific synaptic partners ^36^ and electrophysiological properties ^3,37,38^. It is difficult to distinguish between the subtypes of Purkinje cells and nuclei neurons during *in vivo* recordings in the cerebellum because the different populations only partially follow anatomical boundaries within the cerebellar cortex and nuclei. Additionally, even though the mean firing frequency and pattern are different, the range of these parameters overlap between molecularly distinct subtypes of Purkinje cells and nuclei neurons ^3,35,39^. Therefore, the diversity of firing properties within whole populations of Purkinje cells and nuclei neurons represents inter-cellular variability, provided by the heterogeneity of intrinsic properties and synaptic inputs, plus intra-cellular variability, provided by the dynamic impact of the many sensorimotor inputs. Thus, the complexity and dynamics of cerebellar function offer an inroad to explore data sampling.

Ultimately, changes in the spiking activity of cerebellar neurons have a direct relevance to neurological disorders. In mouse models of cerebellar movement disorders, the source of cerebellar dysfunction is consistent with changes in the firing rate and/or pattern of Purkinje cell and cerebellar nuclei neuron firing ^7,13,40–49^. Altered basal firing properties have also been reported in the thalamus ^11^, basal ganglia ^50^, and cerebral cortex ^51^ during abnormal movement. The task of recording firing activity from neurons in the awake condition has many challenges and limitations, particularly in movement disorders, as the awake animal may frequently perform involuntary actions that can destabilize neural recordings ^52^. Many experiments make use of anesthetized preparations to address these limitations ^28^. However, the presence of the chosen anesthetic and particular anesthesia regime can also affect the firing features of neurons ^53,54^ and anesthesia-induced immobility could limit sensorimotor-related changes in neural firing ^20,21,46,55–57^. Likewise, fluctuations in firing properties associated with the relative arousal state of the animal may be relevant to ongoing behavior ^58–60^. This raises the question, within the constraints of standard experimental settings, how can we ensure that biologically relevant differences are represented in the spiking activity of single neurons when comparisons are made between experimental groups?

In this study, we investigate how experimental constraints on data sampling, such as recording duration and experimental replicates, affect the analysis of *in vivo* neuronal activity. We use two principal neuron types of the cerebellum to examine a diverse set of neuronal spiking properties. We find that recording duration influences the measurement of variability of firing rate over short timescales. Additionally, we find varying degrees of cell-to-cell variability within neural populations. These findings have implications for determining the recording duration and number of cells per animal that are needed to make meaningful comparisons across experimental groups.

## Results

### In vivo spiking activity of cerebellar nuclei neurons fluctuates over time

The basal firing properties of cerebellar nuclei neurons are often represented as values that describe the firing rate and pattern of spike activity. Both firing rate and pattern are determined by the timing of inter-spike intervals ^13,61^. The firing rate, or frequency, is represented by the number of inter-spike intervals sampled over a specific recording duration (spikes/s). Firing pattern is determined by the variation in the timing of inter-spike intervals. Therefore, firing pattern also represents the irregularity of the firing rate that can be described on two different timescales, local and global. The “local irregularity” refers to spike-to-spike variability, or the variability of the instantaneous firing rate, also known as CV2. The “global irregularity” refers to the variability of inter-spike-intervals sampled over a given recording period, or the variability relative to the mean firing rate of the recording duration, also known as the coefficient of variation (CV). Frequency, CV2, and CV are among the most common parameters used to describe basal neural activity and are therefore ideal measures to use when determining the impact of experimental design on the representation of firing properties that are extracted from spike level data.

When investigating the potential differences in spiking activity between disease models or experimental groups, one needs to ensure that the baseline firing properties are optimally represented. This means that the recording of baseline activity must have a duration that encompasses the average state of a neuron during that recording session within the specific recording setup. A common arrangement for this type of recording involves head-fixing a mouse over a rotating/running wheel (Figure 1A, top). There is a large amount of sensorimotor modulation that we expect from this setup as the mouse can move all parts of its body, except for the head, and it sits exposed to the recording room, allowing it to interact with and perceive a range of sights, smells, and sounds. Indeed, we find that the firing rate of cerebellar nuclei neurons fluctuates during our recording sessions (Figure 1B).

We first exemplified how recording duration influences the precision and accuracy of parameter estimations by comparing the deviations of parameter values calculated from shorter and longer recording durations in a small sample set of representative neurons. We predicted that longer recording durations provide more robust estimations of firing parameters, closer to the “true” parameter value. It follows that measurements that are both accurate and precise would be tightly clustered around the measurements from the longest recording. Measurements that are precise, but not accurate, would tightly cluster around a value that deviates from the true parameter value. Measurements that are accurate, but not precise, would center on the true parameter value with deviations on either side.

Overall, when we compared parameter estimations in short recording durations (10s, 20s, 30s) to those in longer recordings (Figure 1B), we found that most measurements of firing rate (frequency), global irregularity (coefficient of variation, CV), and local irregularity (CV2) were precise (open circles in Figure 1C-E). Only select sampling durations had a parameter value that deviated by more than 10% from the value calculated over a longer, 60 s, recording window (closed circles in Figure 1C-E). Measurements of firing frequency were accurate, despite some measurements lacking precision, as over- and underestimations occurred with equal incidence (Figure 1C). In contrast, the majority of global irregularity (CV) measurements were underestimated in the shorter recordings (Figure 1D). This suggests that shorter recording durations may not provide an accurate representation of CV by skewing the measurement to lower values. Finally, we saw relatively small deviations in CV2 estimations between shorter and longer recording durations (Figure 1E), indicating that short recording durations may represent the local firing variability accurately and precisely.

### Shorter duration recordings underestimate the variability in cerebellar nuclei neurons

Next, we set out to investigate whether recording duration influences the accuracy and precision of frequency, CV, and CV2 in most neurons in our database. If the deviation of parameters occurs systematically, lacking accuracy with a consistent valence such as with the underestimation of CV, shorter recordings may not fully represent the natural fluctuations of firing rate within a neuron. To test this, we investigated the differences found between shorter and longer recording lengths in frequency, global irregularity, and local irregularity. We selected nuclei neurons recorded in control mice from which we had a long, stable recording (180 s). From these recordings, we randomly sampled one long recording duration (120 s) for each neuron, which we call “reference recordings.” We also sampled several recording durations of varied length for each neuron (10 s – 120 s, in 10 s intervals), which we call “sample recordings.” We then calculated the difference in frequency, CV, and CV2 between the randomly selected reference and sample recordings. We repeated this process a hundred times, each time selecting a new sample duration from the same recording, for each nuclei neuron (Table 1, n=33).

**Table 1.** Summary statistics for neurons included in our analyses and figures. Values represent mean ± SD. T-tests were used to find statistically significant differences in Frequency, CV, and CV2 of nuclei neurons and Purkinje cells recorded in awake control mice versus anesthetized control mice, or awake control mice versus awake experimental mice. P-values smaller than 0.05 are in bold italics.

| Fig | Neuron Type | Spike Type | n | Awake/<br>Anesthetized | Recording Length (s) | Frequency (Hz) | CV | CV2 |
| --- | --- | --- | --- | --- | --- | --- | --- | --- |
| 1 | Nuclei neuron | All spikes | 33 | Awake | 224.5 ± 44.9 | 67.2 ± 24.1 | 0.49 ± 0.21 | 0.41 ± 0.11 |
| 2 | Purkinje cell | Simple spikes | 16 | Awake | 225.1 ± 39.1 | 66.6 ± 22.9 | 0.50 ± 0.09 | 0.43 ± 0.10 |
| 2 | Purkinje cell | Complex spikes | 16 | Awake | 225.1 ± 39.1 | 1.43 ± 0.39 | 0.74 ± 0.08 | 0.84 ± 0.07 |
| 3 | Nuclei neuron | All spikes | 17 | Anesthetized | 317.6 ± 63.5 | 34.4 ± 12.6<br>*p*<0.001 | 0.38 ± 0.15<br>p=0.051 | 0.39 ± 0.13<br>p=0.586 |
| 3 | Purkinje cell | Simple spikes | 33 | Anesthetized | 304.0 ± 37.4 | 44.4 ± 14.4<br>*p*<0.001 | 0.38 ± 0.10<br>p=0.054 | 0.35 ± 0.10<br>*p*=0.005 |
| 3 | Purkinje cell | Complex spikes | 33 | Anesthetized | 304.0 ± 37.4 | 1.15 ± 0.43<br>*p*=0.035 | 0.75 ± 0.18<br>p=0.675 | 0.90 ± 0.15<br>p=0.143 |
| 4 | Nuclei neuron | All spikes | 16 | Awake | 220.7 ± 86.2 | 54.1 ± 31.0<br>p=0.112 | 0.82 ± 0.27<br>*p*<0.001 | 0.56 ± 0.18<br>*p*=0.001 |
| 4 | Purkinje cell | Simple spikes | 14 | Awake | 215.8 ± 53.3 | 32.4 ± 19.3<br>*p*<0.001 | 1.45 ± 0.66<br>*p*<0.001 | 0.72 ± 0.20<br>*p*<0.001 |

We found that the differences between the reference and sample recordings occurred in either direction for firing frequency (Figure 1F) and local irregularity (CV2; Figure 1H), resulting in a mean difference from the reference recording of near zero for all sample durations. However, the mean global irregularity was systematically lower than the estimate in the reference recording for samples between 10 s and 50 s, which is represented as a negative average of CV difference (Figure 1G). A systematic underestimation of the CV could provide an inaccurate representation of the true global irregularity in firing patterns of the recorded neurons.

We next investigated whether this systematically skewed representation of global variability significantly deviates from the “true” mean of the reference recordings. To test this, we paired one reference recording and one sample recording from each of the neurons (n=33), found the mean reference and sample values of the population, and then performed a paired t-test. This procedure was repeated 100 times to obtain a percentage of statistically significant test results. We then repeated the entire process a total of 25 times, each time with a new reference recording for each neuron, to measure the average percentage of statistically significant test results from the population. Since the pairs of parameter values were calculated from recordings from the same neuron in the same mouse during the same recording session, any statistically significant differences from the reference recording would represent a skewed lack of accuracy in the sample recordings. This is because even if the individual measurements lacked precision, the population mean sampling value should be close to the mean reference value if the measurements are accurate. Finding a statistically significant difference between the reference and sample group suggests that the sample population has trended away from the reference population value. Therefore, an experimenter may have made a different conclusion about the neuron’s properties if the sample recording duration was analyzed instead of the reference duration. We refer to this as a “false positive” conclusion because a significant difference has been found where there should be none. For these reasons, we did not expect the proportion of false positives to be above the level at which we accepted significance (5%). Indeed, the false positive rate was equal to, or smaller than, 5% for all comparisons between frequency and local irregularity (CV2) for recordings from all sample durations included in our analysis (Figure 1I). However, for the global irregularity parameter (CV), this false positive rate was much higher than 5% for sample durations that were shorter than 60 s (Figure 1I).

After finding that sampling durations of less than 60 s resulted in false positive errors, we sought to determine whether this statistically significant difference represented a meaningful magnitude of difference between the populations. We tested this by investigating the percentage of cells that had a large deviation between their parameter estimation from the reference recording and that from their sample recording durations. A large deviation was considered to be an absolute difference of 10% or more between the estimated and reference parameter, anything less than which would be considered an acceptable magnitude of deviation. We also set a threshold of 80% for the minimum acceptable percentage of cells that were free from major deviations. This way our analysis only sought out recording durations resulting in a large percentage of cells with largely deviating measurements. We measured the proportion of comparisons that had an absolute difference of 10% or more, for 100 random sample recordings of each duration. We repeated this comparison for 25 different reference recordings and took the average proportion of neurons that had parameter estimations within a 10% range of the reference recording. We found that for sample durations of 60 s or longer, more than 80% of the neurons had a global irregularity estimation within a 10% range of the reference recordings (CV, Figure 1J).

We acknowledge that 60 s-long recordings might be a challenging goal for many experimental paradigms, especially when recording from awake and moving animals. Therefore, we also investigated how quantification using our shortest recording duration (10 s) compared to parameter quantification over longer recording lengths (Figure 1K). We found a high false positive rate when we compared recording durations of 10 s-long to 30 s or longer recording durations. These findings show that a difference in recording duration is sufficient to demonstrate statistically significant differences in the spiking activity between groups of neurons because the firing rate variability is underestimated in shorter recording durations. If short recordings are used, care must be taken to closely match the recording durations for all cells included in the analysis.

### Shorter duration recordings underestimate the variability in Purkinje cell firing properties

Given that nuclei neuron recordings show a dependency on recording length for an accurate and precise representation of global irregularity, we next set out to investigate whether these observations are also true for a second principal neuron type in the cerebellum, Purkinje cells (Figure 2A). Purkinje cells fire two types of action potentials, simple spikes and complex spikes (Figure 2B) ^62,63^, that differ in origin, shape, and frequency. Simple spikes are spontaneously generated and occur between 40 and 200 Hz *in vivo* in mice (Table 1). Complex spikes are generated by input from climbing fibers originating in the inferior olive, which is located in the brainstem, and occur at a much lower firing frequency (1-2 Hz, Table 1) ^21,63–66^. Both simple spike and complex spike firing frequencies fluctuate during recording sessions in mice (Figure 2B). We analyzed the relationship between recording duration and representation of frequency, CV, and CV2 as described for cerebellar nuclei neurons and only included stable Purkinje cell recordings from control animals with a duration of 180 s or longer (Table 1, n=17).

**Figure 2.**
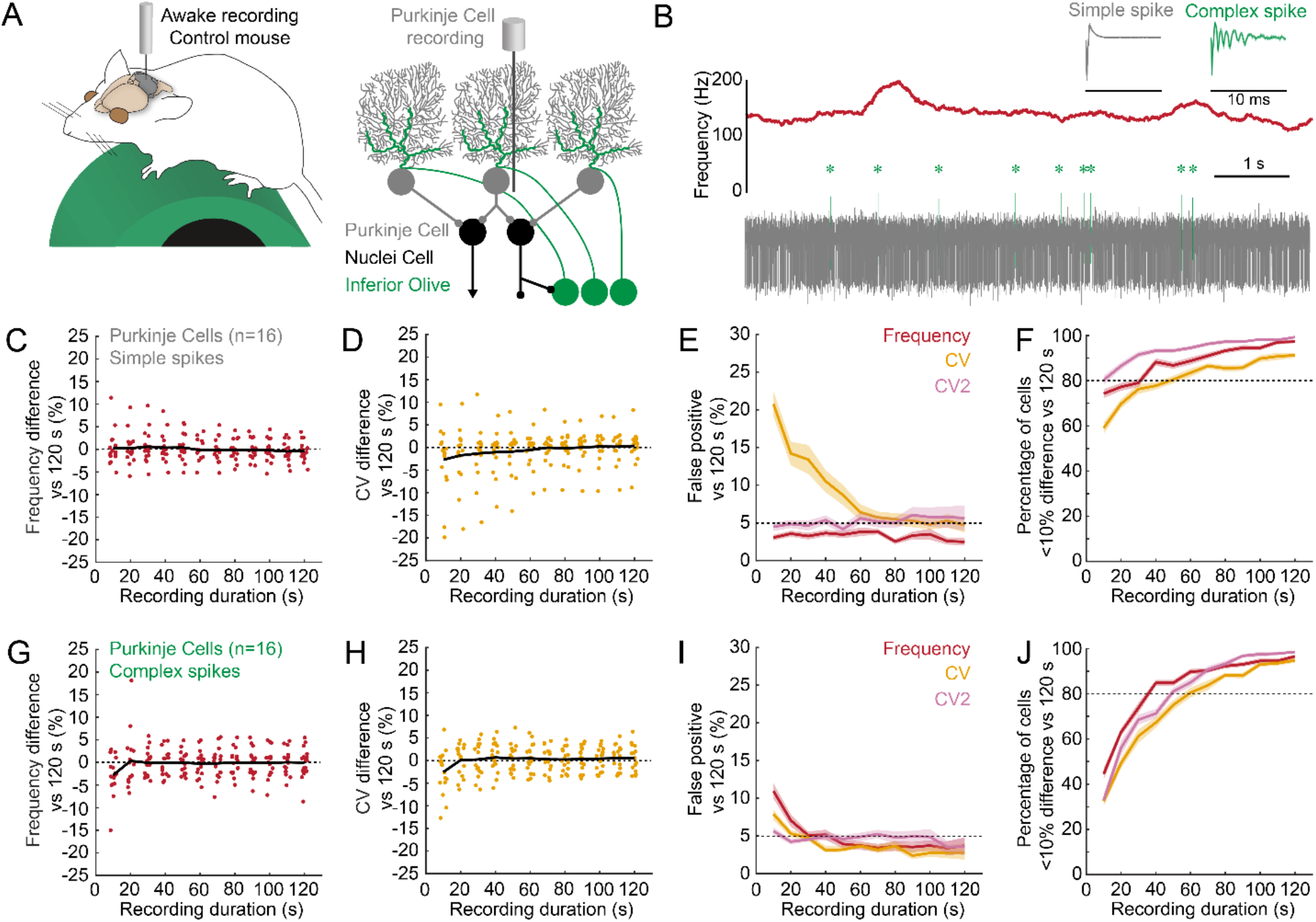
Variability in simple spike and complex spike firing properties of Purkinje cells in awake mice. **A**. Schematic of recording setup depicting an awake mouse on a wheel (left). Schematic of cerebellar circuit with Purkinje cell recording (right). **B**. Example mean frequency (calculated over 0.5 s) trace from a Purkinje cell recording with variability in firing pattern (middle, red). Duration of example trace is 7.5 s. Bottom shows the underlying raw electrophysiological recording trace with simple spikes in grey and complex spikes in green. The higher power view traces (average waveforms across a 20 s segment of a recording, top. 10 ms scale, below) demonstrate the unique spike profiles that distinguish simple spikes from complex spikes. **C**. Frequency and **D**. CV difference from a randomly sampled 120 s-long (reference) recording, extracted from a recording >180 s, calculated from simple spikes. Each dot represents the mean difference of 100 randomly sampled recordings for each recording length for each neuron. The black line represents the average deviation from the frequency or CV calculated in the 120 s-long (reference) recording. The black line diverging from the dotted line at zero in **D**. shows systematic underestimation of CV in shorter recording durations. **E**. Number of significant paired t-tests between 25 randomly sampled 120 s-long (reference) recordings and 100 randomly sampled recordings for each varied recording duration for each neuron, simple spikes only. Mean +/- SEM. Frequency is shown in oxblood red, CV in orange, and CV2 in pink. **F**. Percentage of neurons within 10% deviation from the frequency (oxblood red), CV (orange), and CV2 (pink) of 25 randomly sampled 120 s-long (reference) recordings, each compared to 100 randomly sampled recordings for each varied recording duration for each neuron, simple spikes only. Mean +/- SEM. **G**. Same as **C**., but for complex spikes. **H**. Same as **D**., but for complex spikes. **I**. Same as **E**., but for complex spikes. **J**. Same as **F**., but for complex spikes.

Like the cerebellar nuclei neurons, frequency measurements from Purkinje cell simple spike sample recordings consisting of varied durations (10 – 120 s) deviate from those measured from a reference recording (120 s). However, the average deviation converges to zero even for the sample recordings of the shortest duration (10 s) (Figure 2C). In contrast to firing frequency, global irregularity (CV) is systematically underestimated in shorter sample recordings (shorter than 60 s) (Figure 2D), which leads to a high rate of false positives differences (Figure 2E). Finally, we found that for sample recordings longer than 50 s, each of the Purkinje cell simple spike parameters had crossed the threshold of more than 80% of parameter estimates being within a 10% range of the reference recording parameter estimates (Figure 2F). This shows that Purkinje cell simple spikes also require a recording duration of ∼60 s for accurate and precise descriptions of their firing properties and that short sample durations underestimate the global irregularity in spiking activity.

Next, we investigated whether recording length also influences the ability to estimate the firing properties for complex spikes, which are characterized by their lower frequency. We found that, unlike the higher frequency firing rate in nuclei neurons and Purkinje cell simple spikes, the complex spike frequency was systematically underestimated in sampling recording durations of 10 s (Figure 2G). We also found that, for the complex spikes, the global irregularity was underestimated in sampling recordings with a duration of 10 s (Figure 2H). This resulted in a high rate of false positive differences for sample durations of 10 – 20 s compared to reference recordings for the frequency and CV parameters, but not local firing irregularity CV2 (Figure 2I). Finally, we found that for sample durations of 60 s or longer, parameter estimations of more than 80% of the neurons are within a 10% range of the reference recording (Figure 2J). Remarkably, the sample duration necessary to exceed this 80% threshold is longer for low-frequency complex spikes than the high-frequency Purkinje cell simple spikes and nuclei neurons, showing that precise representation of low-frequency complex spike firing patterns requires a longer recording duration.

### Firing rate variability estimations are less dependent on recording length in anesthetized mice

We hypothesized that the relatively long recording duration (∼60 s) needed to accurately represent the baseline firing patterns relies on capturing sufficient sensorimotor events that modulate the nuclei neuron and Purkinje cell firing rates ^20,21,46,57,67^. If insufficient sensory or motor events are captured in a given time window, the range of firing rates, and therefore the variability, would be underestimated. In anesthetized animals, immobility should prevent motor-event related changes in Purkinje cell and nuclei neuron firing rate, while firing rates are still modulated by sensory stimuli ^53,68–70^. The measurements of firing parameters should be less dependent on sample duration if fewer events lead to fluctuations in cerebellar neuron firing pattern in anesthetized mice. We therefore mined our data to pinpoint the recording duration that would be necessary to accurately represent CV in recordings from anesthetized mice (Figure 3A-C).

**Figure 3.**
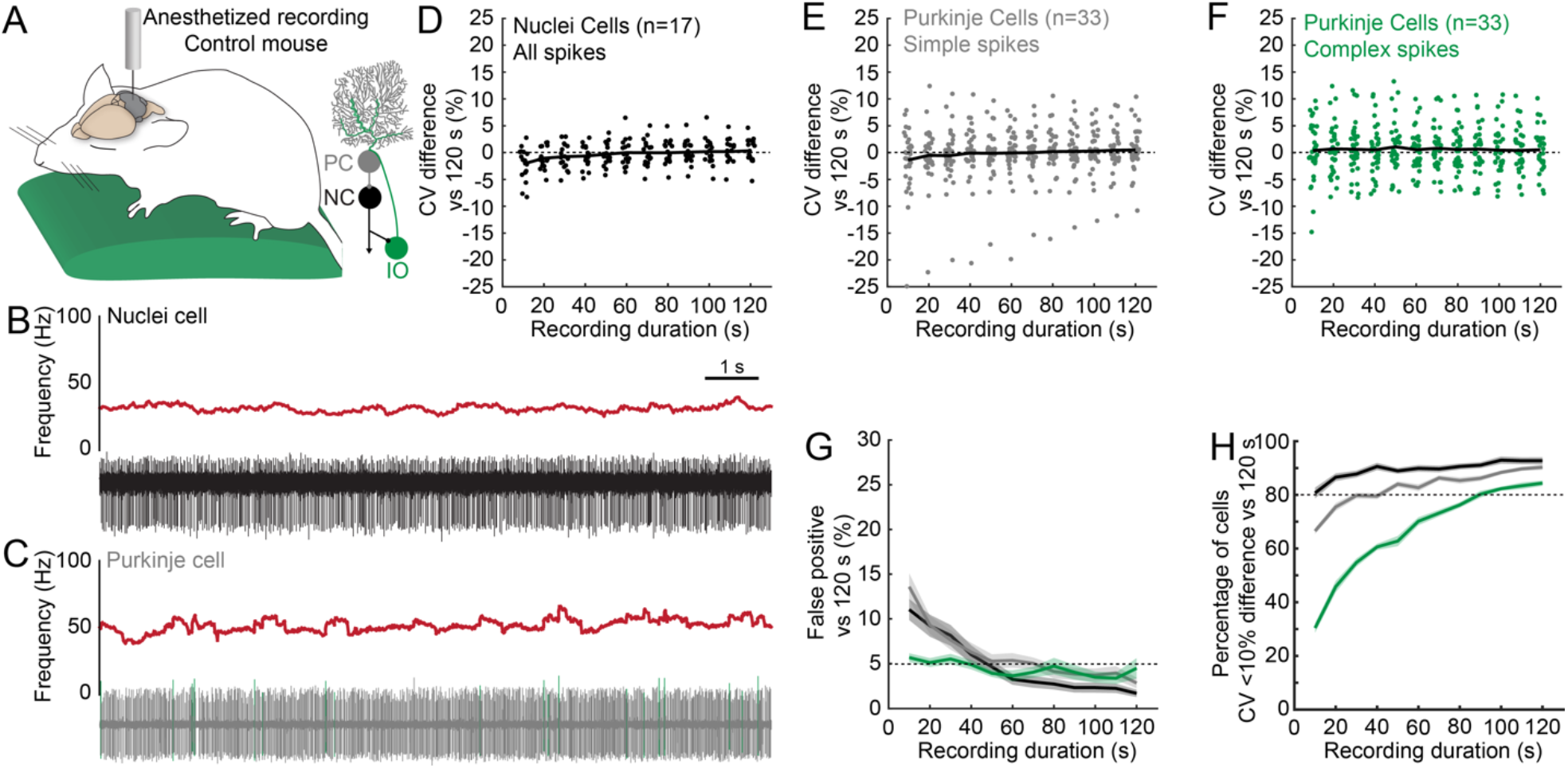
Variability in the firing properties of cerebellar nuclei neurons and Purkinje cells in anesthetized mice. **A**. Schematic of recording setup depicting an anesthetized mouse on a heating pad (left). Simplified schematic of cerebellar circuit (right). **B**. Example trace of a nuclei neuron recording in an anesthetized mouse. Frequency as calculated over 0.5 s intervals (top) and corresponding raw electrophysiological recording (bottom). Example trace is 10 s long. Scale = 1 s. **C**. Example trace of a Purkinje cell recording from an anesthetized mouse. Frequency as calculated over 0.5 s intervals (top) and corresponding raw electrophysiological recording (bottom). Simple spikes are depicted in gray and complex spikes in green. Example trace is 10 s long. Same scale as in **B. D**. CV difference from a randomly sampled 120 s-long (reference) recording duration, extracted from a recording >180 s, for nuclei neurons, all spikes. Each dot represents the mean difference of 100 randomly sampled recordings for each recording length for each neuron. The black line represents the average deviation from the CV calculated in the 120 s-long (reference) recording. **E**. Same as **D**., but for simple spikes in Purkinje cells. **F**. Same as **D**., but for complex spikes in Purkinje cells. **G**. Number of significant differences in CV measured using paired t-tests between 25 randomly sampled 120 s-long (reference) recordings and 100 randomly sampled recordings for each varied recording duration for each neuron, simple spikes only. Mean +/- SEM. Nuclei neurons, all spikes in black. Purkinje cells, simple spikes in gray. Purkinje cells, complex spikes in green. **H**. Percentage of neurons within 10% deviation from CV in 25 randomly sampled 120 s-long (reference) recordings, each compared to 100 randomly sampled recordings for each varied recording duration for each neuron. Mean +/- SEM. Nuclei neurons, all spikes in black. Purkinje cells, simple spikes in gray. Purkinje cells, complex spikes in green.

We found that for cerebellar nuclei neurons in anesthetized mice (Table 1, n=17), the difference in CV compared to a reference recording converges to zero at sample durations of 40 s or longer (Figure 3D). The percentage of false positives between the reference recording and sampling recording converges to 5% in sample durations over 40 s, as well (Figure 3G). However, the relative imprecision of estimation of global irregularity is relatively small, as more than 80% of the CV calculated in the sample durations of all lengths are within a 10% range of the CV estimated in the reference recording (Figure 3H).

We separately analyzed simple spikes and complex spikes from Purkinje cell recordings in anesthetized mice (Table 1, n=33). For simple spikes, we found that the average difference between the sample and reference recordings converged to zero at sample durations over 20 s (Figure 3E) and the percentage of false positives between reference recording and sample recordings converged to 5% for sample recordings over 40 s (Figure 3G). For sample durations of 30 s or longer, more than 80% of neurons had a CV estimate within a 10% range of the CV calculated in the reference recording (Figure 3H).

For complex spikes, we found that the average difference in CV as calculated between the sample and reference recordings was around zero for all sampling durations tested (Figure 3F), and the false positive rate was under 5% for all durations (Figure 3G). Interestingly, less than 80% of the neurons had a CV estimate within a 10% range of the reference recording when the CV was calculated in sample recordings shorter than 90 s. Thus, complex spike CV estimates in shorter sample recordings are imprecise, but do not skew systematically in either direction and therefore do not result in significant differences between the shorter sampling recordings and the long reference recordings.

Taken together, in line with our hypothesis, we find that the recording duration necessary to accurately represent the CV in the cerebellar nuclei neurons and Purkinje cells recorded in anesthetized mice is shorter (∼40 s) than in awake mice (∼60 s). Even though we observed a trend to lower CV in neurons recorded in anesthetized mice compared to neurons recorded in awake mice (Table 1), none of the reductions were statistically significant (nuclei neuron: p=0.051; Purkinje cell, simple spikes: p=0.053; Purkinje cell, complex spikes: p=0.675). This suggests that the observed differences in the recording length to optimally represent the neural variability are not solely due to a reduction in global irregularity and may be secondary to a reduction in environmentally-derived events that modulate cerebellar firing rates in immobile mice.

### Highly variable firing properties are accurately represented in short recordings

Abnormal movements in mouse models of motor disorders may complicate obtaining stable recordings. Our group and others have found a high incidence of an increase in variability in firing rate in mouse models for ataxia, dystonia, and tremor (Figure 4A) ^7,8,71,72^. Therefore, spike variability can arise from the actual stability of the setup itself or from mechanistic changes in the firing properties of disease-associated neurons. We therefore analyzed whether recording duration also influenced the representation of variability in mouse models with unusually high CV. We included neurons from which we obtained a recording of 120 s or longer from mouse models of ataxia, dystonia, and tremor that have high firing rate irregularity in cerebellar nuclei neurons (Figure 4B) and Purkinje cells (Figure 4C) compared to neurons recorded in control mice (Table 1) (nuclei neuron: p<0.001; Purkinje cell, simple spike: p<0.001).

**Figure 4.**
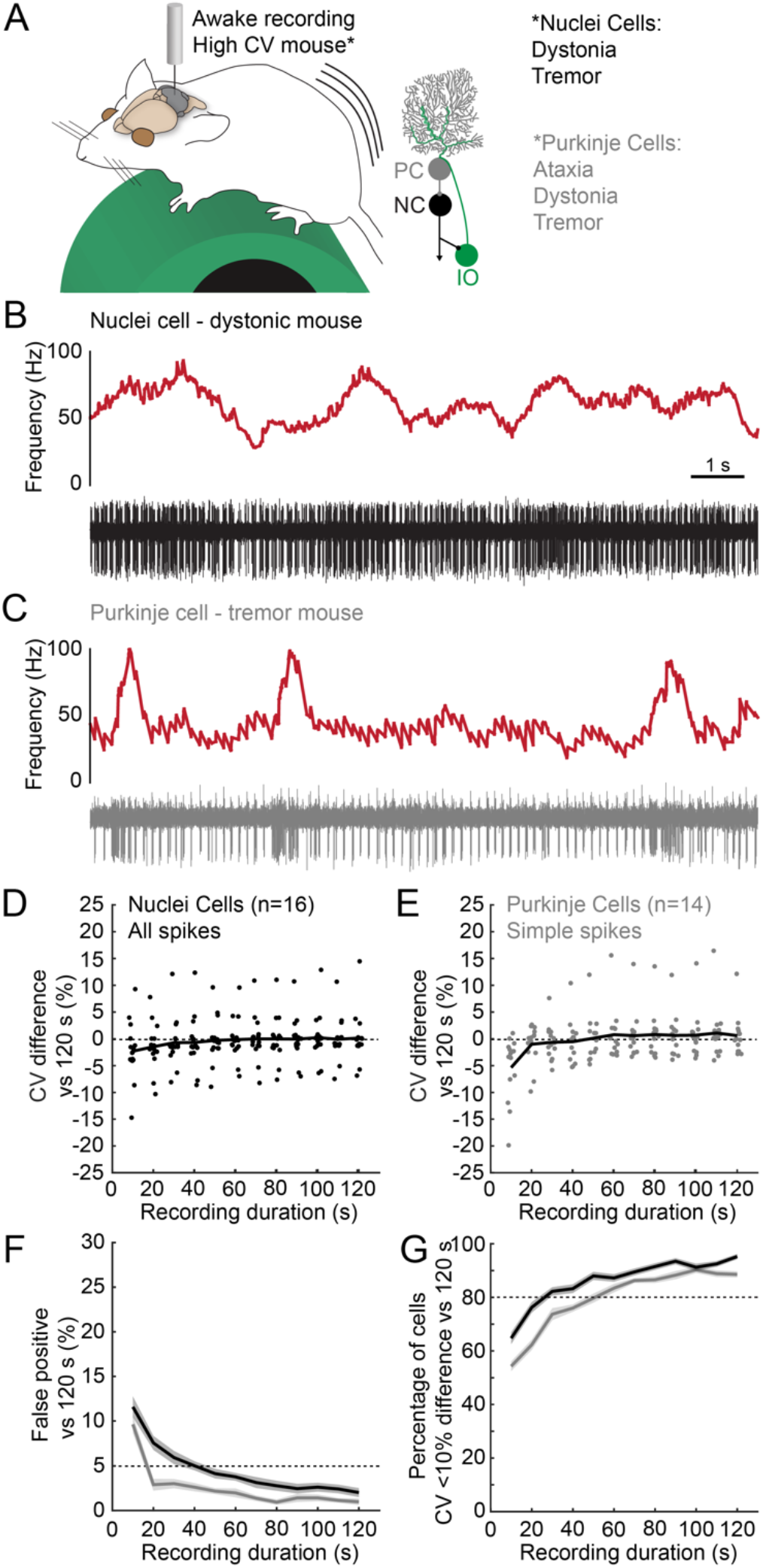
Variability in the firing properties of cerebellar nuclei neurons and Purkinje cells in mouse models of disease. **A**. Schematic of recording setup depicting an awake mouse on a foam running wheel (left). Simplified schematic of cerebellar circuit (right). Neurons included in the analysis for this figure come from *Ptf1a*^*Cre*^*;Vglut2*^*fl/fl*^ (dystonia-like) mice ^8^; harmaline-injected (tremor) mice ^7^; and *L7*^*Cre*^*;Vgat*^*fl/fl*^ (ataxia) mice ^7^. **B**. Example trace of a nuclei neuron recording in an awake dystonic mouse. Frequency as calculated over 0.5 s intervals (top) and matched raw electrophysiological recording (bottom). Example trace is 10 s long. **C**. Example trace of a Purkinje cell recording in an awake tremoring mouse. Frequency as calculated over 0.5 s intervals (top) and matched raw electrophysiological recording (bottom). Example trace is 10 s long. Same scale as in **B. D**. CV difference from a randomly sampled 120 s-long (reference) recording duration from a recording 120 s, for nuclei neurons from disease models, all spikes. Each dot represents the mean difference of 100 randomly sampled recordings for each recording length for each neuron. The black line represents the average deviation from the CV calculated from the 120 s-long (reference) recording. **E**. Same as **D**., but for simple spikes in Purkinje cells. **F**. Number of significant differences in CV measured using paired t-tests between 25 randomly sampled 120 s-long (reference) recordings and 100 randomly sampled recordings for each varied recording duration for each neuron. Mean +/- SEM. Nuclei neurons, all spikes are in black. Purkinje cells, simple spikes are shown in gray. **G**. Percentage of neurons within 10% deviation from CV in 25 randomly sampled 120 s-long (reference) recordings, each compared to 100 randomly sampled recordings for each varied recording duration for each neuron. Mean +/- SEM. Nuclei neurons, all spikes are in black. Purkinje cells, simple spikes are shown in gray.

In the cerebellar nuclei (Table 1, n=16), we observed that at sampling recording lengths of 30 s or longer the average difference between reference recordings and sample recordings converges to zero (Figure 4D), the false positive rate converges to 5% (Figure 4F), and the number of neurons with a CV within a 10% range of the reference recording reaches 80% (Figure 4G). This shows that the CV of nuclei neurons with a highly variable firing pattern are accurately and precisely represented in recording durations of 30 s or longer.

For Purkinje cells in mice with high CV (Table 1, n=14), we only see a systematic underestimation of CV in recordings with a sample duration of 10 s (Figure 4E) and concomitantly only in the recordings with a sample duration of 10 s do we observe false positive differences (Figure 4F). The number of neurons with a CV within a 10% range of the reference recording reaches 80% at sampling recording lengths of 50 s or longer (Figure 4F). Together, this shows that for Purkinje cells with high global irregularity, recording durations as short as 10 s can provide a sufficiently accurate, albeit imprecise, representation of the firing rate variability.

### Within mouse variability does not drive systematic differences in nuclei neuron firing properties

In addition to variability within the firing rate of single neurons, there is heterogeneity in firing properties across the population of recorded neurons. This heterogeneity can arise because of differential responses to sensory information or motor commands and may therefore be dependent on the mouse’s behavior during a specific recording session. As a result, multiple neural recordings from the same mouse may not be true independent samples, but rather nested data ^73^. When we investigate multiple nuclei neurons recorded from the same awake control mouse (Figure 5A), we see differences in the firing frequency and fluctuations in the firing rate (Figure 5B). We hypothesized that if recordings from the same mouse are truly nested data and dependent on the behavioral state of the mouse, the relative difference in firing parameters would be smaller within the same mouse than between different mice. For our analysis, we only included neurons with a recording duration of 60 s or longer and only control mice from which we obtained three or more nuclei recordings that met this criterion. This resulted in a sample of 92 neurons from 21 mice. For each of the 92 neurons we calculated the relative difference in frequency, CV, and CV2 by dividing the absolute difference in parameter estimation for each neuron by the sum of the parameter estimations (see Methods). We visualize these relative differences in frequency (Figure 5C), CV (Figure 5D), and CV2 (Figure 5E) using heatmaps, with blue representing the smallest difference and red representing the largest differences (Figure 5A). The order of neurons is based on the mice from which they were obtained, indicated by the labels on the x- and y-axes. If there was a relatively smaller difference in parameter estimation from neurons obtained from the same mouse, we would expect to see deep blue colors on the top-left to bottom-right diagonal where we plot the differences between neurons obtained from the same animal, and warmer coloring away from the diagonal. Visually, we did not observe this trend in any of the parameters tested (Figure 5C-E).

**Figure 5.**
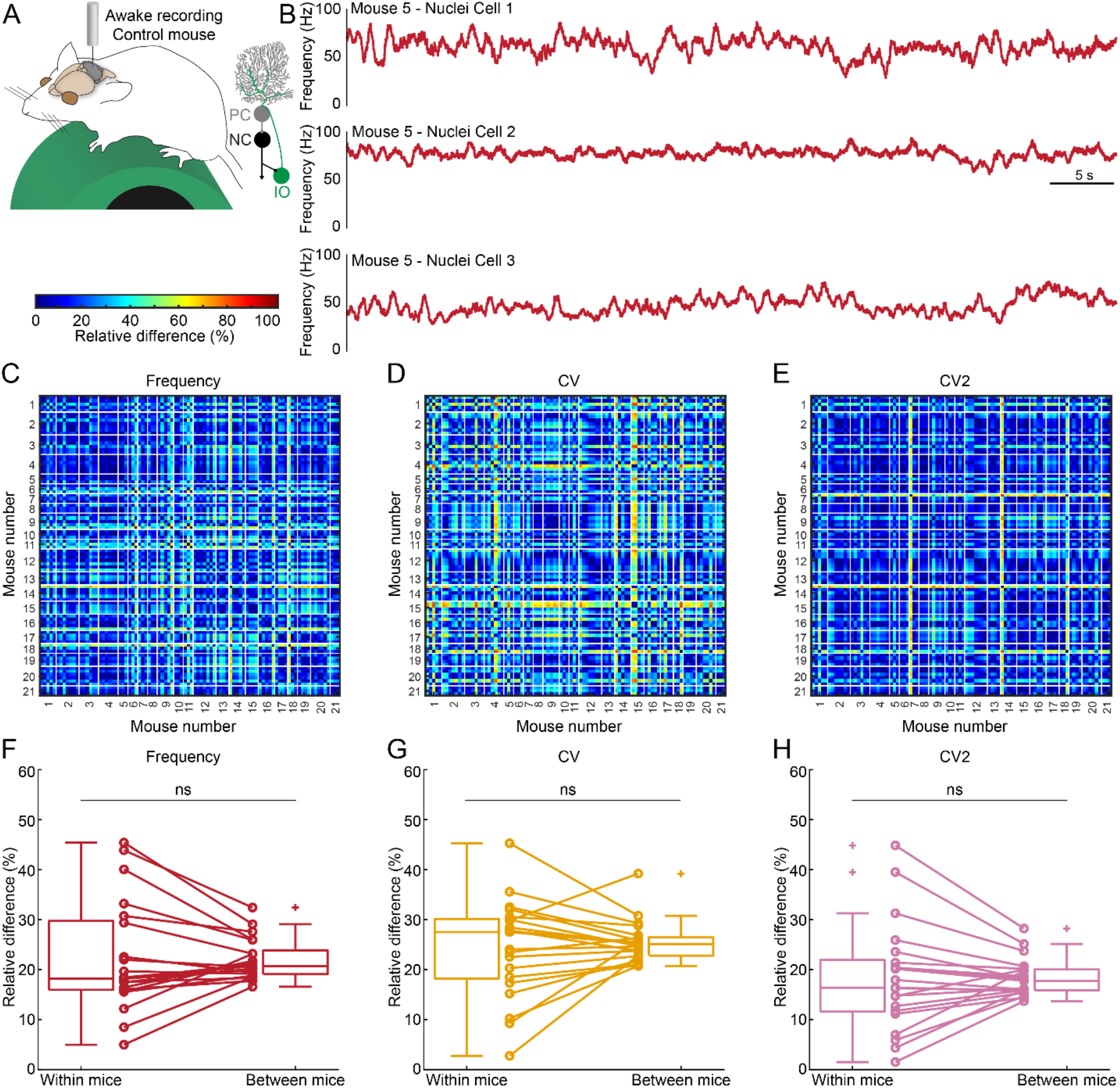
Differences in the firing properties of cerebellar nuclei neurons measured within and between awake mice. **A**. Schematic of recording setup depicting an awake mouse on a foam running wheel (left). Simplified schematic of cerebellar circuit (right). **B**. Example mean frequency traces of nuclei neuron recordings in awake control mice. All three traces come from the same mouse and represent the frequency calculated over 0.5 s intervals. Traces are 30 s long. **C**. Relative difference in frequency between all neuron pairs in 21 mice. Each row and column represent 1 neuron. Thin white lines segregate neurons from different mice. **D**. Same as **C**. although for CV. **E**. Same as **C**. for CV2. **F**. Normalized mean relative percent difference in frequency between nuclei neurons from within the same mice or between mice, statistically analyzed using a paired t-test. Not significant (ns). **G**. Same as **F**. although for CV. **H**. Same as **F**., although for CV2.

Next, we calculated the relative difference in nuclei firing frequency (Figure 5F), CV (Figure 5G), and CV2 (Figure 5H) within each mouse and between other mice in the data set. We do this by first averaging all the relative differences between each pair of neurons obtained in the same mouse (within mice difference) followed by averaging the relative differences between all pairs of neurons in that mouse and all neurons from other mice (between mice difference). We did not find any statistical differences between the relative within mouse differences compared to the relative between mice differences for frequency (p=0.906), CV (p=0.516), or CV2 (p=0.941) (Figure 5 F-H). We also did not observe that the within mouse differences were smaller than the between mouse differences from nuclei neurons recorded in anesthetized mice or mouse models for motor disease (Table 2). Together, these analyses suggest that the sampling of three or more cerebellar nuclei neurons from the same mouse during the same recording session does not bias the parameter estimation of cerebellar nuclei neurons and that individual recordings from nuclei neurons can be confidently analyzed as independent samples that represent the population.

**Table 2.** P-values for differences in firing properties in neurons recorded in experimental groups selected based on their distinct relevance to healthy or diseased circuits. Neurons included are from previously published studies^7,13,28,91^. Significant differences are in bold italics.

| Experimental group | Neuron type | Spike type | Mice (N),<br>neurons (n) | p-value<br>Frequency | p-value<br>CV | p-value<br>CV2 |
| --- | --- | --- | --- | --- | --- | --- |
| Anesthetized<br>Control mice | Nuclei neuron | All spikes | N=4, n=19 | 0.696 | 0.235 | 0.283 |
| Awake<br>Harmaline-injected mice | Nuclei neuron | All spikes | N=3, n=11 | 0.312 | 0.245 | 0.741 |
| Awake<br>*Ptf1a<sup>Cre</sup>;Vglut2<sup>fl/fl</sup>* mice | Nuclei neuron | All spikes | N=4, n=14 | 0.431 | 0.951 | 0.995 |
| Awake<br>*Pcp2<sup>Cre</sup>;Vgat<sup>fl/fl</sup>* mice | Nuclei neuron | All spikes | N=3, n=9 | 0.301 | 0.982 | 0.572 |
| Awake<br>*Thap1<sup>+/-</sup>* mice | Nuclei neuron | All spikes | N=6, n=29 | 0.749 | 0.839 | 0.790 |
| Anesthetized<br>Control mice | Purkinje cell | Simple spikes | N=11, n=46 | 0.445 | 0.452 | 0.197 |
| Anesthetized<br>Control mice | Purkinje cell | Complex spikes | N=11, n=46 | **0.002** | **0.034** | 0.247 |
| Awake<br>*Thap1<sup>+/-</sup>* mice | Purkinje cell | Simple spikes | N=4, n=20 | 0.081 | 0.342 | 0.489 |
| Awake<br>*Thap1<sup>+/-</sup>* mice | Purkinje cell | Complex spikes | N=4, n=20 | 0.345 | 0.574 | 0.905 |

### Variability between mice drives systematic differences in Purkinje cell firing rate

We next set out to investigate whether the lack of mouse-dependent effects was also true for the simple spikes and complex spikes recorded in Purkinje cells in awake mice (Figure 6A). Similar to nuclei neurons, we find that Purkinje cells recorded from the same mice during the same recording session have variable firing properties (Figure 6B). Next, we repeated the calculation of relative differences in frequency (Figure 6C-E), CV (Figure 6G-J), CV2 (Figure 6K-N) for simple spikes and complex spikes using the same methods as we described for the nuclei neurons. Again, we included only neurons from which we obtained a stable recording with a duration of 60 s or more, and only control mice with at least three individual neural recordings. We included a total of 56 Purkinje cell recordings from 13 control, awake mice in our analysis.

**Figure 6.**
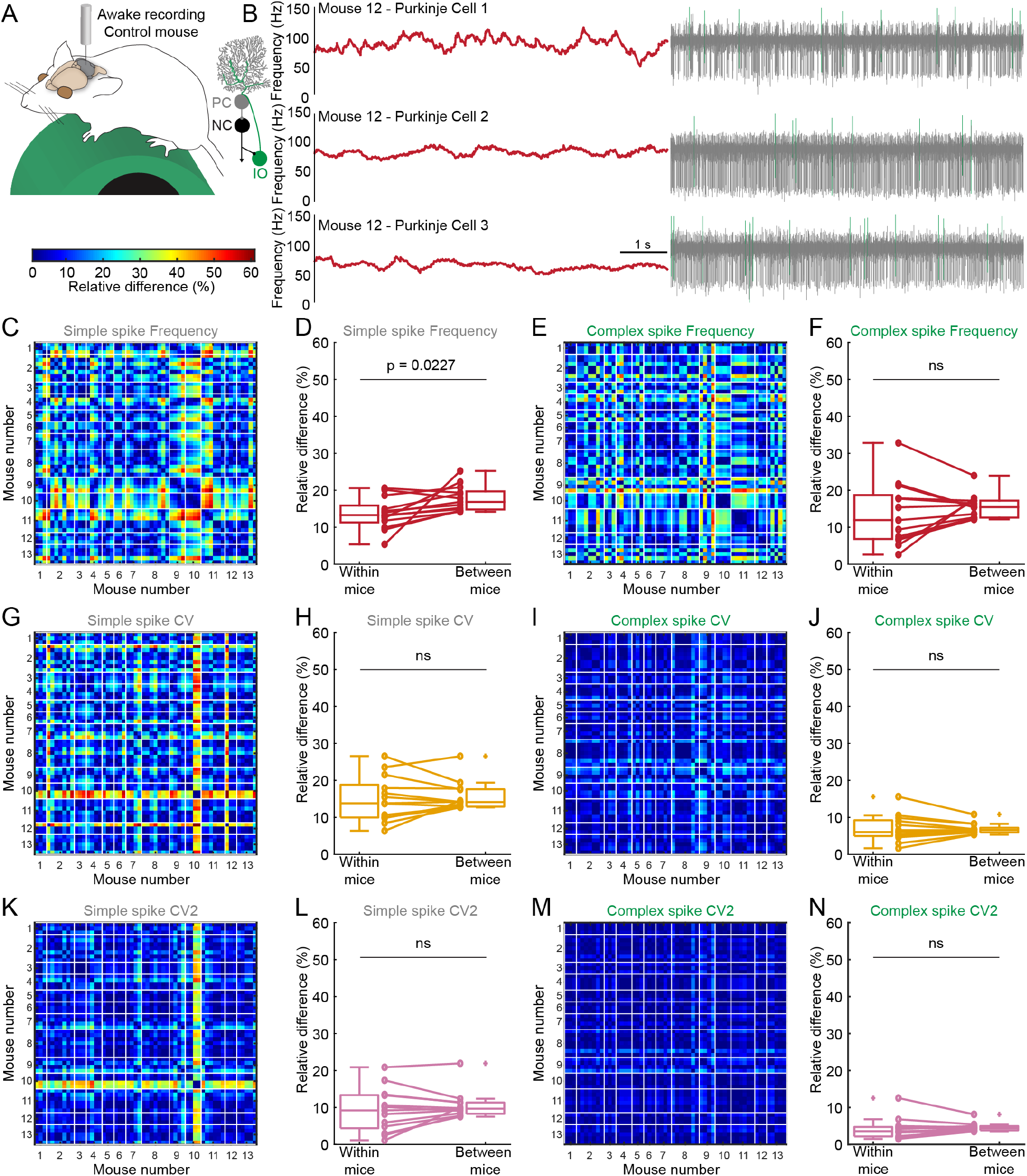
Differences in the firing properties of Purkinje cells measured within and between awake mice. **A**. Schematic of recording setup depicting an awake mouse on a foam running wheel (left). Simplified schematic of cerebellar circuit (right). Heatmap scale (bottom). **B**. Example traces of Purkinje cell recordings in awake control mice. All three traces come from the same mouse and are 7.5 s long. Mean frequency calculated over 0.5 s intervals is represented in red (left). Simple spikes are represented in gray and complex spikes in green (right). **C**. Relative difference in simple spike frequency between all neuron pairs in 13 mice. Each row and column represent 1 neuron. Thin white lines segregate neurons from different mice. **D**. Mean relative difference in frequency of Purkinje cell simple spikes from neurons within the same mice or between mice, statistically analyzed using a paired t-test. **E**. Same as **C**. for complex spike frequency. **F**. Same as **D**. for complex spike frequency. Not significant (ns). **G**. Same as **C**. for simple spike CV. **H**. Same as **D**. for simple spike CV. **I**. Same as **C**. for complex spike CV. **J**. Same as **D**. for complex spike CV. **K**. Same as **C**. for simple spike CV2. **L**. Same as **D**. for simple spike CV2. **M**. Same as **C**. for complex spike CV2. **N**. Same as **D**. for complex spike CV2.

We found a statistically significant higher relative difference in simple spike firing frequency between mice compared to within mice (p=0.0227, Figure 6D). We did not observe any statistically significant effects on the relative difference in complex spike firing frequency between mice compared to within mice (p=0.326). Similarly, we did not find mouse-specific effects on the CV in simple spikes (p=0.560) or complex spikes (p=0.811) or on the CV2 in simple spikes (p=0.250) and complex spikes (p=0.473). When we extended our analysis to Purkinje cells recorded in anesthetized mice, we found that only complex frequency (p=0.002) and CV (p=0.034) were more similar within mice than between mice (Table 2). Moreover, we did not find mouse-specific effects in a mouse model for genetic dystonia (*Thap1*^*+/-*^ mice, Table 2). Thus, under certain conditions, Purkinje cell firing activity may be influenced by mouse-dependent effects and *in vivo* Purkinje cell recordings obtained from the same mouse cannot always be assumed to unequivocally represent independent measurements that properly report on the entire population.

## Discussion

In this study, we investigated how data sampling influences the representation of spiking activity in cerebellar neurons. We first set out to investigate how recording duration influences the estimation of firing parameters in cerebellar nuclei neurons, and simple and complex spikes in Purkinje cells. We found that recording duration influences the ability to appreciate a neuron’s full repertoire of firing patterns. Specifically, shorter recordings resulted in a skew towards lower estimates of global variability in all neuron types, mouse models, and experimental paradigms included in our analysis. Thus, recording duration itself affects the representation of basal firing properties and recording length should be accounted for when investigating the firing parameters in recording sessions *in vivo*. In the recordings included in our analyses, we find that 60 s-long recordings are sufficient to avoid systematic underrepresentation of firing rate variability, but this duration may be longer, or shorter, for other brain regions, species, or experimental settings. Regardless of the experiment, we recommend matching the recording duration used to extract estimations of parameters describing firing properties for all neurons included in a given experiment to avoid statistical significance arising from a difference in recording length rather than the inherent firing properties in diverse populations of neurons.

Next, we set out to investigate whether neural recordings obtained from the same animal are independent of each other or are susceptible to mouse-dependent variations. We found that for nuclei neurons in our database, there are no mouse-dependent effects on firing properties in awake or anesthetized control, or disease-model mice. In contrast, Purkinje cell simple spike firing rate may be more similar in neurons recorded from the same mouse than in neurons recorded from different mice. Finally, in the anesthetized state, we observe that Purkinje cell complex spike firing properties are more similar within mice than between mice, which may result from minor differences in anesthesia levels. Together, these results show that presentation and analyses of Purkinje cell firing properties may have to take mouse-specific effects in account. This could be accomplished by showing data in super plots ^74^ and performing statistical analyses that take inter-mouse variability into account and corrects for nested data ^73,75^. These results also underscore that data from multiple neurons recorded in multiple mice should be included when changes in Purkinje cell firing rates are anticipated and reported. Ideally, similar numbers of neurons should be included for each mouse used in the analyses.

At first glance, it may be surprising that there seem to be mouse-dependent variability in Purkinje cell firing properties, but not in nuclei neuron firing properties, as Purkinje cells provide direct inhibitory input onto nuclei neurons. However, there are several cerebellar properties that may drive mouse effects only in Purkinje cells and not nuclei neurons. First, different Purkinje cell types are organized in longitudinal zones whereas the cerebellar nuclei neuron sub-types are intermingled ^14,76^, even though they also respect the functional modules. The Purkinje cells within these zones have been found to have different baseline firing rates in a spatially organized manner ^1,30,39^. Thus, a single penetration of the electrode through the cerebellar cortex and nuclei is more likely to encounter the same molecular class of Purkinje cells with relatively similar firing rates, and more likely to encounter different molecularly defined classes of nuclei neurons with relatively varied firing rates. Therefore, when recording from neuron types or regions with known spatially organized diversity, the experimenter must either consistently sample neurons of the same molecular or regional identity or consistently sample across identities. Additionally, even though Purkinje cells directly innervate nuclei neurons, the firing rate of nuclei neurons adapts relatively fast to changes in Purkinje cell firing rates ^77^. The lack of a direct inverse correlation between Purkinje cell firing rate and nuclei firing rate is further corroborated by our own findings in mice in which neurotransmitter release from Purkinje cells is prohibited and we do not find a systematic increase in nuclei neuron firing rate ^7,28^. Thus, if mouse-dependent differences in Purkinje cell firing rates are due to differences in motor activity, sensory input, or arousal state during the recording session, nuclei neurons may adapt quickly and not show mouse-dependent effects.

A previous study found that in *in vitro* electrophysiological recordings of pyramidal neurons in the sensorimotor cortex of rats, short observation length results in systematic underestimation of measures of local irregularity ^78,79^. In that analysis, Nawrot and colleagues argued that this is the result of relatively long inter-spike-intervals, which are the primary driver of local irregularity, becoming overlapped with the boundaries of the recording window, which results in these longer inter-spike-intervals being poorly represented in the analysis. A similar argument my also explain the relative underestimation of frequency in the low-frequency complex spikes, wherein the window excludes complex spike rates that can be higher than the typical 1 Hz, even if these events are relatively rare (Figure 2G). However, we find that recording durations of 20 s are sufficiently long to reduce this bias, which are much shorter than the recording durations that include 50-100 inter-spike-intervals that Nawrot and colleagues estimate to be sufficient to eliminate systematic bias in their studies.

Nevertheless, our finding that short recording lengths result in an underestimation of global firing rate irregularity is in line with the mathematical and empirical examples provided in these previous papers ^78,79^. The reason for this might be similar to the effect of a super short recording length (millisecond to seconds) on underestimation of local irregularities, namely, insufficient sampling of periods with relatively high variability. As stated previously, recordings during baseline conditions are not devoid of neurons that are computing signals for ongoing natural tasks. Even head-fixed mice can perform many movements including walking on the wheel, paw, and whisker movements, all of which can directly modulate firing properties in cerebellar neurons ^19,20,80,81^. Similarly, recordings in awake and anesthetized mice conducted in laboratory settings with a myriad of olfactory, visual, auditory, and somatosensory stimuli, which can change the firing rate in cerebellar neurons as well ^21,69,70^. Finally, depending on the context cerebellar neurons may toggle between up- and down-states which can also contributes to firing rate variability ^53,82^. The *in vivo* variability of firing rate in neurons in the cerebellum is likely the culmination of changes in firing rate due to movement, sensation, and inner-cellular state. A longer recording duration is necessary to capture the broader range of firing rates at which the neuron fires in the experimental setting. When a longer duration is not possible, but for that matter is more ideal in all cases, it is best to match recording durations so that a similar sample of neuromodulatory events may be captured in the recordings across neurons, mice, and experimental groups.

Firing rate variability is not unique to the cerebellar nuclei that are modulated by sensory and motor cues, but is also found in motor and sensory cortices and even upon the presentation of the same movement or sensory cue ^83–86^. Many studies have focused on understanding task-specific variability ^87,88^. During task-specific variability, the irregularity in firing rate can be mapped to a specific time-window during which a trained cue is presented. However, when measuring variability in firing rates in lieu of a sensory or motor task, it is unclear what drives the change in firing rate. It is therefore important to adequately sample the firing properties of a neuron over a period long enough to encompass a range of movements, sensory inputs, and internal states.

One particular feature of our study is that we performed our analyses on real neurons recorded *in vivo*, rather than *in silico* modelled spike trains. Our results are therefore independent of model assumptions and reflect robust biological ranges in firing properties. While prior theoretical papers have warned of the effects of nested data in electrophysiological recordings based on modelled assumptions of inter-animal effects ^75^, we show using experimentally obtained data that such effects can indeed occur *in vivo*. Recording duration (or number of spikes) is often selected as an inclusion criterion for analyses on firing properties, yet, to our knowledge, the influence of recording duration has not previously been reported by *in vivo* or *in silico* models of basal spike trains. Nevertheless, we believe that the effect of recording duration is a robust experimental entity that has major relevance to a wide range of neural populations, as we observe that short recordings result in an underestimation of firing rate variability in all spike types, neuron types, experimental settings, and mouse models included in our analyses.

Taken together, while our analyses are focused on variability in firing properties in cerebellar neurons, they may generalize to the representation of firing properties in other neuron types and brain regions as well. We have detailed a set of analyses that provide a straightforward pipeline that scientists can adapt to investigate potential effects of recording duration or mouse-dependent effects on the reporting of firing properties of single neurons. For neural types in which little is known about how recording duration or inter-subject variability influences firing properties, the most conservative approach would be to match recording durations between all experimental groups, record from multiple individuals, include a similar number of recordings per individual, and perform statistical analyses that take inter-subject effects into account.

## Methods

### Animals

All experiments described in this manuscript were reviewed and approved by the Institutional Animal Care and Use Committee (IACUC) at Baylor College of Medicine (BCM). Mice were housed in a Level 3, AALAS-certified facility on a 14-10 light-dark cycle. We used standard breeding paradigms to generate the different alleles, with the date a copulation plug was observed designated as embryonic day 0.5 and the date of birth as postnatal day 0. We used mice from both sexes and a range of ages (two to ten months old). Animals included in the analyses are control mice from mixed genetic backgrounds unless otherwise stated. The majority of neurons included in our analyses were reported in our previous publications (nuclei neurons and Purkinje cell recordings in awake mice: ^7,8,13,89,90^; nuclei neurons and Purkinje cell recordings in anesthetized mice: ^28,91^). Procedures for electrophysiologically identifying different neurons, and collecting the data from different mice, cohorts and preparations by different experimenters was consistent between studies. The several previously reported experimental models are specified in the text. Mice indicated as “dystonic mice” are previously characterized *Ptf1a*^*Cre*^*;Vglut2*^*fl/fl*^ mice ^8^ in which we genetically silence the climbing fiber to Purkinje cell synapse using *Ptf1a*^*Cre*^-mediated ^92^ deletion of a *LoxP*-flanked sequence for the vesicular transporter for glutamate (*Vglut2*) ^93,94^. Mice indicated as “ataxic mice” are previously characterized *Pcp2*^*Cre*^*;Vgat*^*fl/fl*^ mice ^7,28^ in which we genetically silence Purkinje cells using *Pcp2*^*Cre*^-mediated ^95^ deletion of a *LoxP*-flanked sequence for the vesicular transporter for GABA (*Vgat*) ^96^. Tremor mice are previously characterized mice which received a 30 mg/kg intraperitoneal injection of harmaline (Sigma-Aldrich, #H1392) ^7^. *Thap1*^*+/-*^ mice are a genetic model for heredity dystonia that display a mild pathophysiological tremor, which we characterized in a previous report ^13^. For our genetic mouse models, we collected ear-clippings at pre-weaning ages in order to extract DNA for PCR-genotyping. The PCR primers and PCR cycling conditions have all been reported in the aforementioned publications.

### Surgery for recordings in awake mice

We performed a head-plate surgery for awake recordings as described in detail in previous publications ^6–8,52^. Throughout the surgery we kept the mice on a heated surgery pad and stabilized the head using ear bars in a stereotaxic surgery rig (David Kopf Instruments). We provided preemptive analgesics (slow-release buprenorphine at 1 mg/kg subcutaneous or buprenorphine at 0.6mg/kg as well as slow-release meloxicam at 4mg/kg or meloxicam at 5 mg/kg subcutaneous) and maintained mice under continuous anesthesia using nasally delivered isoflurane gas (2-3%). We removed fur from the surgery site and made an incision in the skin over the anterior part of the head. Next, we mounted a custom head-plate over bregma using C and B Metabond Adhesive Luting Cement (Parkell) and placed vertical visible wiring to annotate bregma during recordings. We then used a dental drill to make a circular craniotomy with a diameter of approximately 2 mm over the cerebellum (6.4 mm anterior and 1.3 lateral from Bregma) and placed a custom 3D-printed chamber over the craniotomy that we filled with antibiotic ointment prior to closing. We firmly attached the 3D-printed chamber and headplate to the mouse skull using Metabond and dental cement (dental cement powder #525000; solution #526000; A-M Systems). After surgery, the mice recovered from anesthesia in a clean cage resting on a heating platform. We monitored mice for stress and pain for a minimum of three days after surgery and throughout the experimental period. We provided mice with additional meloxicam and buprenorphine for three days after surgery.

### Surgery for recordings in anesthetized mice

We performed our surgeries in anesthetized mice as described in detail in our previous publications ^5,8^. However, some brief points to note are as follows. Throughout the surgery and recording, we kept the mice on a heated surgery pad and stabilized the head using ear bars in a stereotaxic surgery rig (David Kopf Instruments). Prior to surgery, we anesthetized mice using a mixture of intraperitoneal administered ketamine and dexmedetomidine and nasally-delivered isoflurane gas (1-2%). Next, we removed fur from the head and made an incision in the skin over the anterior part of the head. We used a dental drill to make a craniotomy with a diameter of approximately 5 mm over the cerebellum (6.4 mm anterior and 1.3 lateral from Bregma). Penetrations of the cerebellum proceeded as to capture the activity of as many Purkinje cells and cerebellar nuclei neurons as possible per recording session, which typically lasted for approximately 2-3 hours.

### In vivo electrophysiological recordings

We stabilized the mouse’s head using the surgically attached head-plate (awake mice) or ear bars (anesthetized mice), and kept the mouse on a freely rotating foam wheel (awake mice) or heating pad (anesthetized mice) for the durations of the recording session. We used tungsten electrodes with an impedance of ∼8 MΩ for our recordings. The movement of the electrodes was controlled using a motorized micromanipulator (MP-225; Sutter Instrument Co). The electrical signals obtained by the electrodes were amplified and bandpass filtered (0.3-13 kHz) (ELC-03XS amplifier, NPI Electronic Instruments) before being digitized (CED Power 1401, CED). All *in vivo* electrophysiology signals were recorded and analyzed using Spike2 software (CED).

### Analysis of in vivo electrophysiological recordings

All single neuron recordings were quality controlled with signal to noise ratio as a primary consideration and spike sorted in Spike2. We also considered cell-specific features. We accepted all neurons that had clearly distinguishable complex spikes (a single large spike followed by three to five smaller amplitude spikelets, with a pause in simple spike activity last ∼ 20-50 ms following this whole signature), usually obtained between 0.5 and 2.5 mm from the brain surface as Purkinje cells. We identified the cerebellar nuclei neurons based on the depth at which they were recorded (between 2.5 and 3.5 mm from the brain surface) and their generally lower firing rate compared to Purkinje cells. We only committed to analyzing the full duration of recordings that maintained a stable action potential amplitude.

### Calculation of parameters describing neural firing properties

We described the firing properties of single Purkinje and cerebellar nuclei neuron recordings using a combination of three parameters that are most often reported when describing the firing properties of cerebellar neurons ^2,13,97^. First, we calculated the firing frequency as follows: 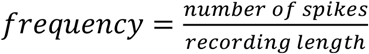. Second, we measured the global irregularity in firing rate, CV, based on the interspike interval (ISI) between spikes: 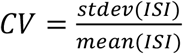. The CV is a standardized measure for the global irregularity in the firing rate as it is calculated based on the average deviation of ISI from the mean ISI (ISI^-1^ is frequency) and normalized to the mean ISI. As a result, a neuron with a highly fluctuating firing rate, will have a high CV, whereas a tonically firing neuron will have a low CV. Third, we calculated the CV2 as follows: 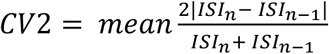 This measure indicates the local irregularity in the firing rate of a neuron as it takes in account the average difference in ISI between each adjacent pair of spikes in the recording ^97^. All analyses were performed using custom-written code in MATLAB (The MathWorks Inc., version 2021a).

### Quantification of the influence of recording length on firing pattern parameters

We determined whether recording length biases the description of firing properties in single neuron recordings by calculating how the frequency, CV, and CV2 differed when calculated over shorter versus longer periods taken from the same recording. In this calculation, we only included neurons from which we had a high-quality, continuous recording of three minutes minimum (180 s). We then randomly sampled a continuous 120 s period from this longer recording and calculated the frequency, CV, and CV2 in this period as an estimation of the true firing properties during our recording session. We choose to calculate the firing properties based on a randomly sampled 120 s long recording time window instead of using the complete recording length in order to equalize the duration length across neurons, to minimize the potential confound of electrode movement that is most likely to occur at the start or end of each recording, and to prevent the convergence of parameter calculations being caused by a higher degree of temporal overlap in the part of the recording that was analyzed.

We next calculated the difference between the parameters of our 120 s recordings with those obtained calculated over time windows *t* from 10 s to 120 s at 10 s intervals (10 s, 20 s, 30 s, … 120 s). These time windows were also randomly sampled from the entire duration of the recording we had available for each neuron. We sampled each shorter time window at least 100 times and calculated the difference as follows: 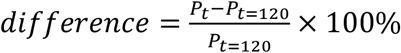. Here, *P* is the calculated parameter (frequency, CV, or CV2) calculated over period of length *t*. The average difference between 100 randomly sampled durations and one longer duration is represented in the figures that show the differences in parameter value.

For our estimation of false positive differences and proportion of neurons that is within 10 % of the reference recording durations, we repeated our analysis 25 times, each with a different randomly sampled 120 s long recording duration. We then performed a paired t-test between the parameter as calculated over recording time *t* and a pre-set recording time (*t=*120 s or *t=*10 s) and counted the number of tests with a p-value lower than 0.05 and for each of the 25 repetitions. Similarly, we counted the proportion of neurons with parameter calculations from the sample recording of duration time *t* that were within 10% of the parameter calculation from the 120 s long recording duration, for each of the 25 repetitions. For these two calculations, the data shown in the figure represents the mean ± the standard error from the mean.

### Quantification of the inter-versus intra-mouse variability in neural firing properties

For our estimation of effect of inter-versus intra-mouse variability, we only included neurons for which we obtained an original recording length of 60 s or longer. We also only included mice for which we obtained at least three neurons that met this criterion. We determined the relative difference in parameter values in neurons as follows: 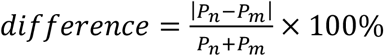. Here, *P* is the calculated parameter (frequency, CV, or CV2) calculated for each neuron *n* and *m*. We then calculated the average relative difference for all neuron pairs where neuron *n* and *m* were obtained from the same mouse (but not the same neuron). We designated this as the relative difference within mice, for each mouse. We also calculated the average relative difference for neuron pairs where neuron *n* and *m* were not obtained from the same animal. We designated this as the relative difference between mice, for each mouse. We investigated whether the deviation between the relative difference within and between mice was statistically significant using a paired t-test.

## Acknowledgements

This work was supported by Baylor College of Medicine (BCM), Texas Children’s Hospital, The Hamill Foundation, and the National Institutes of Neurological Disorders and Stroke (NINDS), R01NS100874 and R01NS119301 to RVS. The study was also supported by the Eunice Kennedy Shriver National Institute of Child Health & Human Development of the National Institutes of Health under Award Number P50HD103555 for use of the Cell & Tissue Pathogenesis Core (BCM IDDRC). The content is solely the responsibility of the authors and does not necessarily represent the official views of the National Institutes of Health. Support was also provided by a Dystonia Medical Research Foundation (DMRF) grant to RVS and a DMRF postdoctoral award to MEvdH.

## Notes

### Competing Interest Statement

The authors have declared no competing interest.

